# Dynamic chromatin accessibility landscape changes following interleukin-1 stimulation

**DOI:** 10.1101/2020.01.29.924878

**Authors:** Matt J. Barter, Kathleen Cheung, Julia Falk, Andreas C. Panagiotopoulos, Caitlin Cosimini, Siobhan O’Brien, Karina Teja-Putri, Graham Neill, David J. Deehan, David A. Young

## Abstract

Genome-wide methods for examining chromatin modification provide detailed information on regulatory regions of the genome. Dynamic modifications of chromatin allow rapid access of the gene regulatory machinery to condensed genomic regions facilitating subsequent gene expression. Inflammatory cytokine stimulation of cells can cause rapid gene expression changes through direct signalling pathway-mediated transcription factor activation and regulatory element binding.

Here we used the Assay for Transposase Accessible Chromatin with high-throughput sequencing (ATAC-seq) to assess regions of the genome that are differentially accessible following treatment of cells with interleukin-1 (IL-1). We identified 126,483 open chromatin regions, with 241 regions significantly differentially accessible following stimulation, with 64 and 177 more or less accessible, respectively. These differentially accessible regions predominantly correspond to regions of the genome marked as enhancers. Motif searching identified an overrepresentation of a number of transcription factors, most notably RelA in the regions becoming more accessible, with analysis of ChIP-seq data confirmed RelA binding to these regions. A significant correlation in differential chromatin accessibility and gene expression was also observed.

Functionality in regulating gene expression was confirmed using CRISPR/Cas9 genome-editing to delete regions for that became more accessible following stimulation in the genes *MMP13*, *IKBKE* and *C1QTNF1*. These same regions were also accessible for activation using a dCas9-transcriptional activator and showed enhancer activity in a cellular model.

Together, these data describe and functionally validate a number of dynamically accessible chromatin regions involved in inflammatory signalling.

## Introduction

DNA is packaged into the nucleus by hierarchical folding around histone proteins to form nucleosomes, followed by further compaction of nucleosomes into chromatin. A dynamic epigenetic code, which constitutes the histone code of histone modifications and nucleosome positioning, as well as other modifications such as DNA methylation, functions to allow, or to prevent, access by transcriptional regulatory machinery to this chromatin [1].

Though DNA methylation is the most well characterised epigenetic mark, changes in gene expression correlate poorly with methylation alterations at promoters, and instead methylation changes tend to occur in regions of the genome defined as transcriptional enhancers [2]. Furthermore, although methyl groups at CpG loci can be both added and removed, the processes involved are not very dynamic, especially active demethylation which requires a series of enzymatic modifications to 5-methylcytosine (5mC) by so-called TET-TDG pathway coupled with base excision repair (BER) [3]. Such pathways are believed to be important for embryonic stem cell differentiation, somatic cell reprogramming and other biological processes involving cell fate transitions, rather than upon rapid activation by cytokines or growth factors.

Dynamic modifications of chromatin can rapidly allow access of condensed genomic regions to gene regulatory machinery. These chromatin modifications can occur either via covalent histone modifying enzymes, such as histone acetyltransferases or methyltransferases, or by ATP-dependent chromatin remodelling complexes, such as SWI/SNF, which are capable of altering the position of nucleosomes along DNA [4, 5]. Genome-wide methods for examining chromatin modification (e.g. ChIP-seq) or chromatin accessibility (e.g. DNase-Seq) have provided detailed information on regulatory regions of the genome however, although powerful, such techniques often require millions of cells and complex, time-consuming, sample preparation. Recently, the Assay for Transposase Accessible Chromatin with high-throughput sequencing (ATAC-seq) method has gained popularity due to its simplicity and low cell number requirement. The assay uses a modified transposase (Tn5) to efficiently cleave exposed DNA while simultaneously ligating sequence-specific adapters. Adapter-ligated DNA fragments are then isolated, amplified by PCR and subjected to next generation sequencing [6].

The pro-inflammatory cytokine interleukin-1 (IL-1) plays a critical role in host responses to pathogens and other harmful agents. To activate cellular responses IL-1 binds to its cognate receptor, IL1R1, which heterodimerises with the common co-receptor IL1RAcP. This tertiary complex then recruits intracellular signalling molecules, including myeloid differentiation factor 88 (MyD88), IL1R-associated kinase (IRAK) and TNF receptor-associated factor 6 (TRAF6) to elicit its cellular response via activation of nuclear factor-κB (NF-κB), as well as mitogen-activated protein kinases (MAPK) p38, c-Jun N-terminal kinase (JNK) and extracellular signal-regulated kinase (ERK) pathways. Given the importance of interleukin signalling a number of clinical syndromes are genetically-associated with a dysregulation of the pathway, while IL-1-related cytokine inhibitors are clinically relevant, especially for a number of auto-inflammatory disorders [7].

Here, we used ATAC-seq to establish dynamic changes in chromatin accessibility following short-term IL-1 stimulation. We identified 241 peaks which are significantly differentially accessible by the cytokine treatment and these are enriched distal from promoters, at ENCODE/Roadmap defined enhancer regions. As anticipated genomic regions that become accessible during the stimulation are significantly enriched for NF-κB (RelA) binding motifs and correlate with gene expression changes following stimulation. We confirm the function of several of these putative enhancers in regulating IL-1-dependent gene expression following CRISPR-Cas9-mediated genomic deletion. Similarly, using a dCas9-transcriptional activator (VPR), gRNAs to these regions could potentiate IL-1-induced target gene expression. Finally, we tested the response of these enhancers to IL-1 in vitro using luciferase reporters. Together, these data define a number of novel, dynamic, inflammatory signalling-dependent enhancer loci.

## Materials and Methods

### Cell culture

SW1353 chondrosarcoma cells (ATCC, LGC Standards, Teddington, UK) were cultured as previously described [8]. As with SW1353, HEK 293T cells were cultured in DMEM. HEK 293T were transfected using FugeneHD (Promega, Southampton, UK) with plasmids lentiCRISPR v2 or lenti-EF1a-dCas9-VPR-Puro, plus packaging plasmids psPAX2 and pCMV-VSV-G (Addgene plasmids #52961, #99373, #12260, and # 8454, respectively). After 48 and 72 hr lentivirus was harvested by centrifugation. SW1353 cells were lentivirus transduced and puromycin-selected to generate SW1353-Cas9 cells, stably expressing Cas9, or SW1353-VPR, expressing dCas9-VPR.

### ATAC-seq

SW1353-Cas9 cells (30,000/cm^2^) were seeded in a 6-well plate, and the next day serum-starved overnight, followed by IL-1α (1 ng/ml) stimulation for 6 hours (each condition in triplicate). Using the Buenrostro *et al*. [6, 9] protocol modified to reduce mitochondrial DNA contamination [10], nuclei were isolated by lysis of cells using ice-cold lysis buffer (10 mM Tris-HCl, pH 7.4, 10 mM NaCl, 3 mM MgCl2 and 0.1% IGEPAL CA-630) and centrifugation at 500*g* for 10 minutes at 4°C. Following isolation, the nuclei were resuspended in the transposase reaction mix (25 μL 2x TD buffer, 2.5 μL Transposase TDE1 (Nextera Tn5 Transposase, Illumina, Cambridge, UK) in a total volume of 50 μL and incubated for 30 minutes at 37°C. All samples were subsequently purified using Qiagen MinElute PCR Purification Kit (Qiagen, Manchester, UK), eluting in 10μl of Elution Buffer. Transposed DNA was PCR amplified using the NEB Next High-Fidelity PCR Master Mix (New England Biolabs, Hitchin, UK) and primer Ad1_noMX in combination with one of six barcoded primers (Table 1) in a 50 μL reaction. Oligonucleotide primer sequences are listed in Supplementary Table 1. Cycling conditions were are previously described [9]. An aliquot of the reaction was subjected to quantitative real-time PCR cycles, the remaining 45 μL was then subjected to PCR for a calculated additional number of cycles as described to avoid amplification to saturation. Finally, the amplified libraries were purified using a Qiagen MinElute PCR Purification Kit and eluted in 20 μl Elution Buffer before quantitation on an Agilent 4200 TapeStation (Agilent Technologies LDA UK Limited, Stockport, UK). Pooled libraries were sequenced on a High-Output Illumina NextSeq500 generating paired-end 75 bp reads.

### Data analysis and Bioinformatics

All samples were sequenced to a depth of 53-74 million paired-end reads (Supplementary Table 2). Quality control of raw sequencing reads was performed using FastQC (v0.11.7). All samples passed QC. Reads were aligned to the hg38 reference genome using Bowtie2 short read aligner (v2.3.4.1) [11] using default settings. Peaks were called using MACS2 (v2.1.1.20160309) [12] with the–nomodel option on. Peaks with an FDR < 0.05 were considered significant. The Integrative Genomics Viewer (IGV) [13] was used to visualise read coverage and peaks. Peaks were annotated using the annotatePeaks.pl program within HOMER (v4.9.1) [14]. The DiffBind (v2.12.0) Bioconductor package was used to find differentially open peaks between IL1 and control samples. For differential peak testing, a consensus peak set was generated by DiffBind using default settings; peaks were considered a consensus peak if it appears in two or more samples. A total of 126,483 unique consensus peaks were found. The DESeq2 method implemented within DiffBind was used to test differential peaks between conditions. Using this method, 241 peaks with an FDR <0.05 were found (Supplementary Table 3). To determine the chromatin state of the peaks, peaks were intersected with chromatin state data from Roadmap E049 cells – bone marrow-derived mesenchymal stem cell (BM-MSC) differentiated chondrocytes [15]. From these cells the 18 state model was collapsed to the 5 representative states and the size of the peak-intersected states summed. Statistical significance of peak enrichment in chromatin states was determined using a Pearson’s Chi-squared test with Yates’ continuity correction. MEME (v5.0.3) [16] was used for *de novo* motif searching in up- and down-regulated peaks, with a minimum motif length of 6bp and a maximum motif length of 30bp. The TOMTOM motif matching tool within the MEME suite of tools was used match motifs found by MEME to known motifs. Genomic Regions Enrichment of Annotations Tool (GREAT) [17] was used to find gene ontology terms related to up- and down-regulated peaks (defined as ≥1.5 fold, p ≤ 0.01), which gave 661 and 1649 peaks, respectively.

### Transcriptome analysis

An Illumina HT-12 v4 microarray was used to measure gene expression in IL-1α treated SW1353 samples (n = 3) compared to controls (n = 3). The lumi Bioconductor package [18] was used to preprocess and normalise the data. Probes with a detection p-value > 0.01 were removed from the analysis. Microarray signals were log2 transformed and normalised using the Robust Spline Normalisation (RSN) procedure within lumi. Differential gene expression analysis was performed using the limma package [19]. A linear model was fitted to the data and an empirical Bayes method implemented within limma was used to find differentially expressed genes between conditions. As data were collected in two experimental batches, this information was added as a covariate in the linear model. Multiple tests were corrected for using the Benjamini-Hochberg method and genes with an FDR < 0.05 were considered to be significant (Supplementary Table 4). ATAC-seq peaks were linked to the transcription start site (TSS) of the nearest gene present on the gene expression microarray using a custom R script.

### Genome-editing

SW1353-Cas9 cells seeded into 6 well plates at 10,000 cells/cm^2^ were transfected with 50μM of each gRNA (two gRNAs per region to delete) duplexed with 100μM tracrRNA using DharmaFECT 1 transfection reagent (Horizon Discovery, Cambridge, UK) following a modified version of the IDT ‘Alt-R CRISPR/Cas9 System’ protocol. The control transfection contained DharmaFECT 1 but no duplex. Cells were incubated at 37°C and after 3 days trypsinised and reseeded into a 75 cm^2^ flask with a small proportion used for genomic DNA isolation with an Invitrogen PureLink Genomic DNA mini kit following manufacturer’s instructions (ThermoFisher Scientific, Life Technologies Ltd, Paisley, UK). Genomic regions of interest were amplified from 100 ng isolated gRNA using Phire Polymerase II (ThermoFisher Scientific) following the manufacturer’s protocol.

### Luciferase analysis

Genomic regions (and deletions) corresponding to the ATAC-seq putative enhancer peaks 21603, 48601, 8848, and 90201 were amplified with Phire II polymerase (ThermoFisher Scientific) with template gDNA used from the corresponding wild-type or Cas9-edited SW1353 cells. PCR oligonucleotides contained additional sequences complementary to *Bgl* II restriction endonuclease digested pGL3-promoter plasmid (Promega, Southampton, UK) for subsequently cloned using the InFusion HD Cloning Kit protocol (Takara Bio, Saint-Germain-en-Laye, France). All plasmids were confirmed by Sanger sequencing. For luciferase analysis, SW1353 cells were seeded into 96 well plates at 18,000 cells/cm^2^. Each well was transfected with 25ng of the corresponding putative enhancer-cloned pGL3-promoter luciferase reporter along with 1.5ng Renilla (pRL-TK Vector, Promega) control reporter vector using FuGENE HD transfection reagent as previously described [20]. Transfected cells were serum starved overnight prior to stimulation with IL-1α (0.5ng/ml) for either 6 hrs or 24hrs as indicated. PBS-washed cells were lysed with passive lysis buffer (Promega) and luminescence monitored using a Glomax Luminometer and the Dual-Luciferase Reporter Assay System (Promega). Firefly luciferase data were normalised to the Renilla luciferase control. An NF-κB-luc reporter (Takara Bio) was used as a positive control (data not shown).

### Data availability

The following published datasets were used in this study. For KB cell RELA ChIP-seq NCBI GEO submission GSE52469 [21, 22], KB/HeLa cell IL-1 ATAC-seq accession GSE134436 [23]. Chromatin state data for Roadmap E049 BM-MSC differentiated chondrocytes accession GSE19465 [15, 24]. ATAC-seq data and microarray data generated here are available with the accession GSE144303 and TBC, respectively.

## Results

### Interleukin-1 signalling initiates dynamic chromatin accessibility changes

To investigate dynamic chromatin signatures during IL-1 stimulation we used the chondrosarcoma cell line SW1353, which has a well-characterised robust response to IL-1 stimulation [25]. A modified ATAC-seq protocol [10] was performed in triplicate on control or IL-1 stimulated cells, generating ≥62 million reads/sample with a low level of contaminating mitochondrial DNA reads (between 1 and 11%) (Supplementary Table 1). The expected nucleosome banding patterns were identified by analysis of fragment size distribution (Supplementary Figure 1A). Interpretation of nucleosome positioning highlighted nucleosome depletion at transcription start sites (TSS) and the position of the most proximal nucleosomes (Supplementary Figure 1B).

126,483 accessible chromatin regions (peaks) were identified across the samples. The genomic distribution of the peaks were annotated with Genbank features to indicate their genic or intergenic localisation (Supplementary Figure 2). Differential accessibility was assessed by comparison of peaks between control and IL-1-treated samples. 241 significant peaks were identified with 64 and 177 regions becoming more or less accessible, respectively, following IL-1 treatment (Figure 1A). Examples of regions that became more or less accessible are given for peaks within the STX8 and FAM69A gene loci respectively (Figure 1B). The majority of the 126,483 peaks are centred around the TSS however, the 241 significant peaks showed an extended distribution with a proportion of regions displaying chromatin changes distal to promoter regions (Figure 1C). Next, we examined peak distribution in relation to chromatin state information obtained from the BM-MSC-derived chondrocytes by the NIH Roadmap Epigenomics project [15, 24]. When assessing all the accessible regions there was increased representation at TSS and enhancer regions compared to the genome-wide distribution of the chromatin states (Figure 1D). For the 241 significantly differentially accessible regions there was a highly significant enrichment at regions marked as enhancers, compared to ‘all peaks’ (*p*<2.2 × 10^−16^) or the genome-wide distribution (*p*<2.2 × 10^−16^) of the chromatin state (Figure 1D). A recent study also identified IL-1 regulated genomic regions in HeLa-derived KB cells [23]. When similarly analysing both datasets, we found 159 more and 137 and less accessible features in common following IL-1 stimulation (Supplementary Figure 3). Of interest, only the more accessible regions following stimulation contained a highly significant overlap (*p*<0.0001) between the datasets, indicating that common genomic regions become accessible following IL-1 stimulation independent of cell-type. Regions that become less accessible could be less cell-type specific or may simply reflect the differences in the initial chromatin states of the diverse cells.

**Figure 1.**
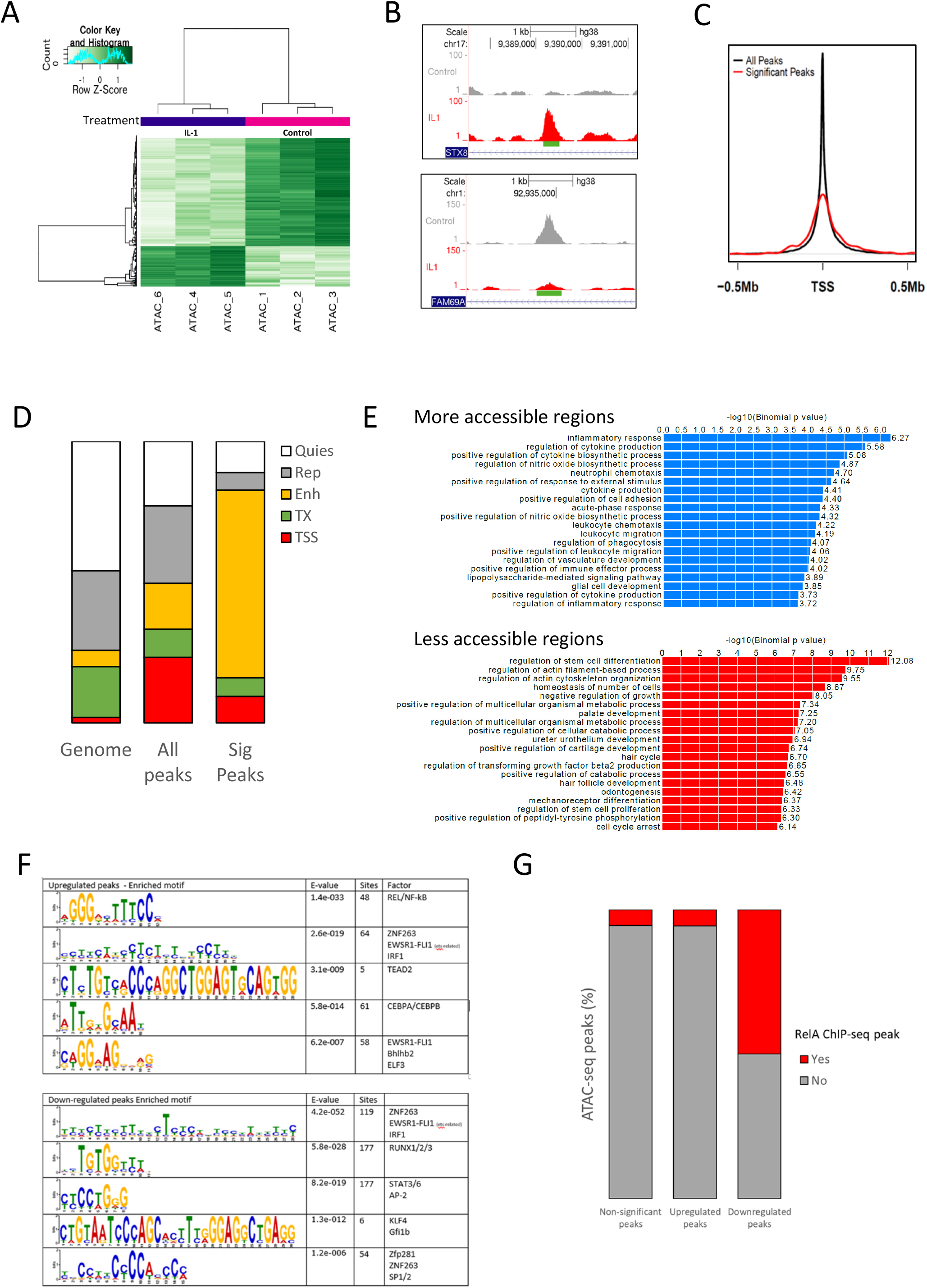
ATAC-seq identified peaks in IL-1-stimulated chondrocytes. SW1353 cells were stimulated in triplicate with IL-1 for 6 hours prior to nuclei isolation and ATAC-seq. A. Heatmap and hierarchical clustering representation of the 241 differentially accessible peaks. B. Genomic vignette of significant peaks within the STX8 and FAM69A gene loci. C. All peaks and significant peak distribution relative to TSS. D. Enrichment of peaks at Roadmap E049 MSC-derived chondrocyte chromatin states. E. GO analysis of more and less accessible peak proximal genes. F. Motif analysis of more and less accessible peaks assessed using MEME and TOMTOM. G. Overlap of more and less accessible peak regions with ChIP-seq-identified IL-1-induced RelA binding sites.

### IL-1-driven accessible chromatin regions are associated with RelA/NF-κB binding

By associating each significant peak with its nearest gene, gene ontology (GO) analysis revealed more accessible regions were associated with terms connected to an inflammatory response or a response to external stimuli (Figure 1E). Terms associated with less accessible regions were more diverse, but interestingly, contained terms associated with cartilage including ‘palate development’ and ‘positive regulation of cartilage development’. Motif analysis to identify potential transcription factors regulating differentially accessible regions indicated regions with increased accessibility were enriched for proteins such as NF-κB (RelA), IRF1/ZNF263, CEBPA, and ELF3 while motifs in regions with reduced accessibility following IL-1 treatment included those for ZNF263, RUNX1 and STAT3 (Figure 1F). Next we intersected the location of our peaks with a RelA ChIP-seq dataset generated following 1 hour IL-1 stimulation of HeLa-derived KB cells [21, 22]. For all our peaks, or the peaks becoming less accessible, only approximately 5% were associated with a RelA peak. In contrast, 50% of the significantly more accessible regions following stimulation contained at least one associated RelA-bound ChIP-seq peak (*p*<2.2 × 10^−16^)(Figure 1G).

### Dynamic alterations in chromatin accessibility correlate with gene expression changes

In order to establish whether the changes in chromatin accessibility were associated with gene expression changes SW1353 cells were again stimulated with IL-1 for 6 hours and whole-genome gene expression microarray analysis performed. IL-1 stimulation resulted in 400 upregulated and 62 downregulated genes (1.5 fold, FDR <0.05) (Figure 2A). Next, we correlated these differentially expressed genes with the changes in chromatin accessibility. The majority of chromatin accessibility changes occurred without an expression change to the nearest gene. However, where gene expression changes were associated with accessibility changes there was a positive correlation between accessibility and altered gene expression (Spearman’s rank correlation rho 0.4771, *p*<1 x10^−7^), suggesting that the opening/closing of chromatin contributes to the observed transcriptional regulation (Figure 2B).

**Figure 2.**
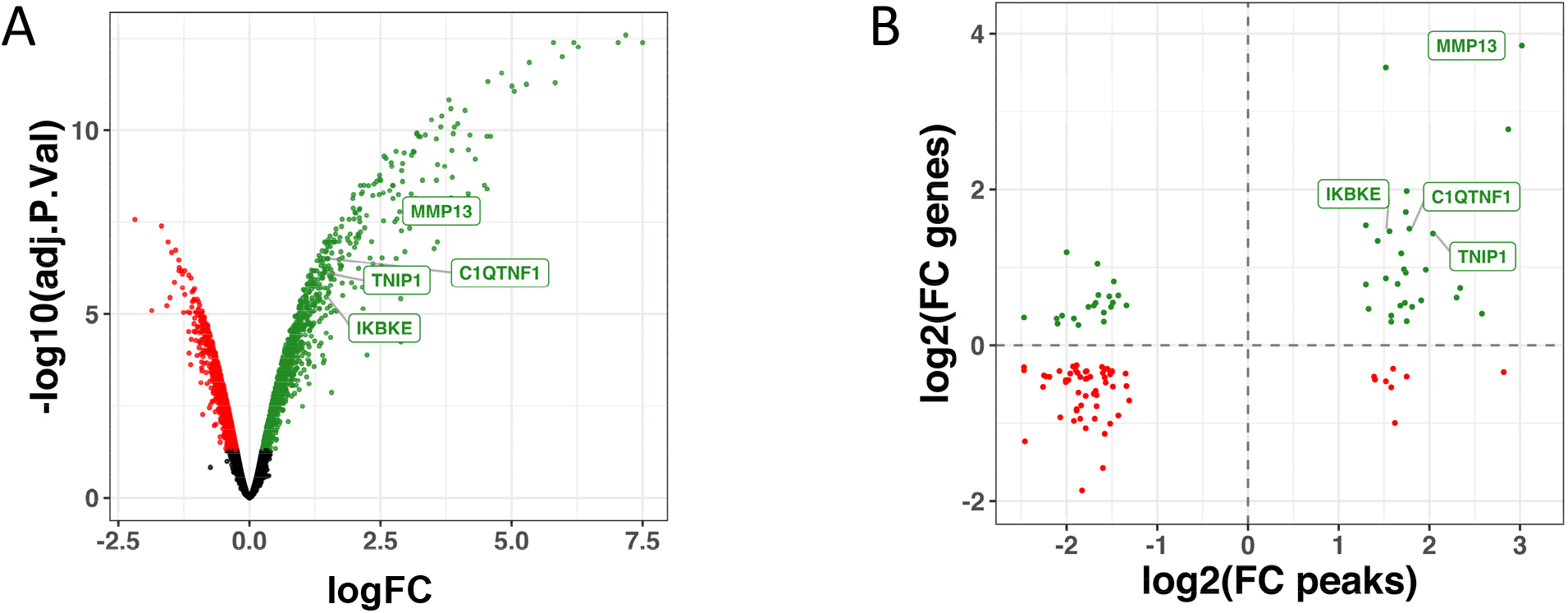
Correlation between differentially accessible chromatin regions and gene expression changes. A. Volcano plot of gene expression from SW1353 stimulated with IL-1 for 6 hours prior to RNA extraction and microarray analysis. Significantly up- and down-regulated genes are represented by green and red dots respectively. B. Comparison of significant ATAC-seq peak accessibility fold change with expression change in proximal gene. Genes labelled were those selected for functional validation.

### Newly accessible regions following IL-1 stimulation can act as enhancers

To test whether regions of increased accessibility at IL-1-induced gene loci were required for gene induction we used pairs of gRNAs and Cas9 to delete the regions encompassing the significant intronic peaks within the *MMP13* (peak 21603), *C1QTNF1* (peak 48601), *IKBKE* (peak 8848) and *TNIP1* (peak 90201) loci (Figure 3A and B). All four genes were upregulated following IL-1 stimulation (Figure 2A). RelA binding sites were identified at these regions by both MEME motif analysis and the intersection with IL-1-induced RelA ChIP-seq peaks (Figure 3A) [21]. Further, interrogation of published ChIP-seq data at the Gene Transcription Regulation Database also indicated the presence of RelA binding sites (Supplementary Figure 5) [26]. We also used JEME (joint effect of multiple enhancers), which predicts enhancer-target gene interactions by leveraging enhancer features and gene expression levels from consortia datasets, to determine the target genes of all our accessible peaks [27]. These predicted interactions for the peaks within *MMP13*, *C1QTNF1*, and *TNIP1* with their respective genes promoter/TSS (Supplementary Figure 6). Cas9-mediated deletion of the regions encompassing the induced-ATAC-seq accessible regions at the *MMP13*, *C1QTNF1* and *IKBKE* loci caused a corresponding reduction in the upregulation of gene expression by IL-1 (Figure 3C). *TNIP1* basal or induced expression was unaffected. Expression of the well-characterised IL-1 responsive gene *IL-8* was unaffected by deletion of any of the four regions, confirming they did not affect NF-κB signalling per se (Supplementary Figure 7).

**Figure 3.**
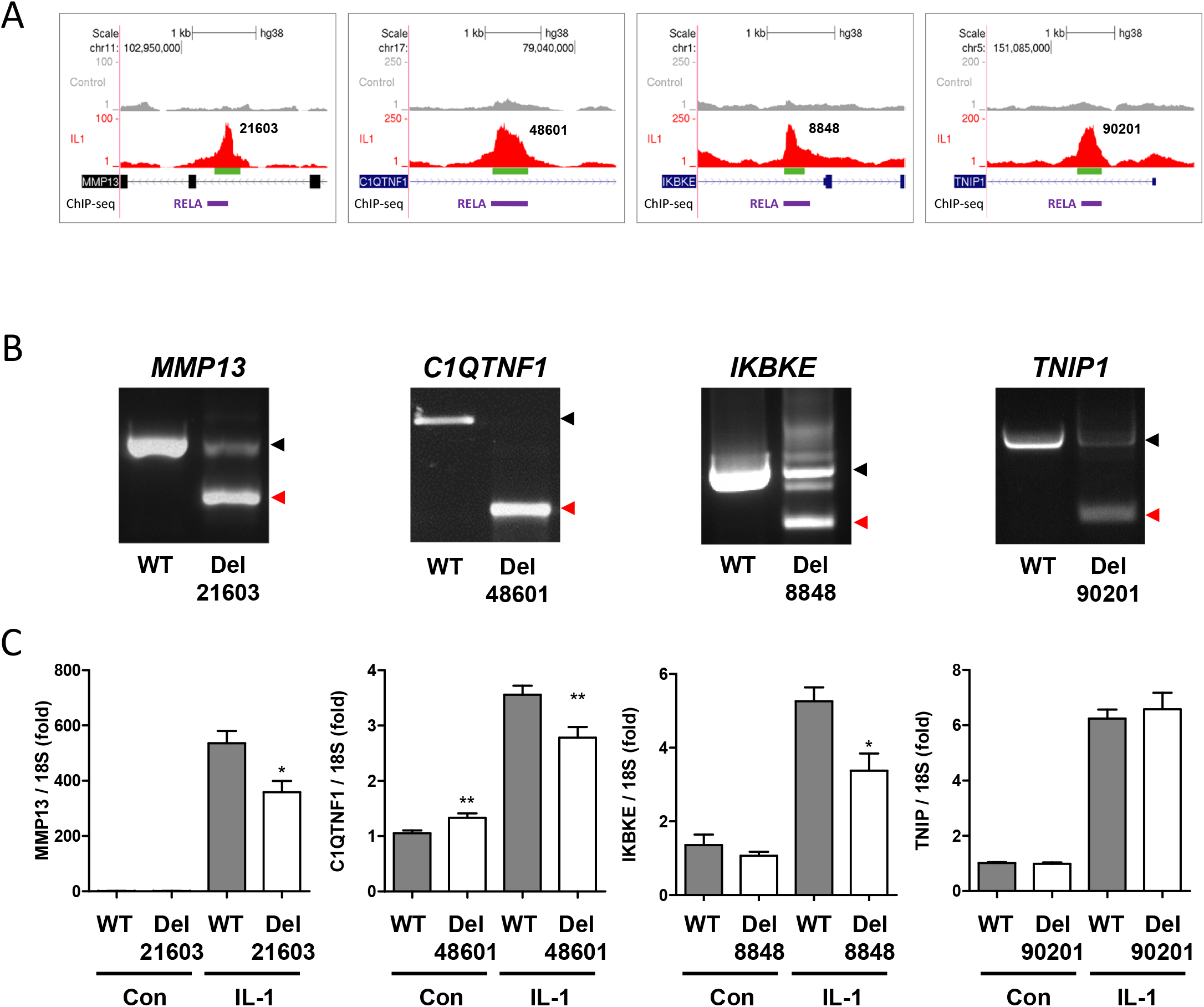
Effect of Cas9-mediated putative enhancer deletion on IL-1-induced gene expression. A. Genomic vignette of significant ATAC-seq peaks within the *MMP13*, *C1QTNF1*, *IKBKE* and *TNIP1* loci following IL-1 stimulation. Numbers 21603, 48601, 8848, 90201 refer to the peak identifier defined by MACs. ChIP-seq-identified IL-1-induced RelA binding sites are depicted below. B. PCR determination of gRNA-targetted Cas9 deletion of ATAC-seq peak regions. Black and red arrrowheads indicate expected wild-type and deletion products, respectively. C. Expression of the indicated genes measured by real-time qRT-PCR in SW1353-Cas9 cells following Cas9-mediated deletion of ATAC-seq regions. Values are mean ± standard error of the mean (SEM) of data pooled from between 3 and 6 independent experiments (each performed in sextuplate). *P<0.05; **P<0.01 for deletion versus wildtype for each control or IL-1 treatment group. Significant differences between sample groups were assessed by a two-tailed Student’s t-test.

The requirement of these regions for IL-1-induced gene expression suggests they act as enhancers. Accordingly, we targeting an activating dCas9-VPR to the ATAC-seq-identified accessible regions to provide further evidence that these putative enhancers regulate local gene expression. Consistent with this, the treatment of SW1353 constitutively expressing dCas9-VPR with gRNAs targeting these regions resulted in upregulation of basal gene expression of *MMP13*, *C1QTNF1* and *IKBKE* (Figure 4). The IL-1-induced expression of all four genes was also increased following gRNA addition.

**Figure 4.**
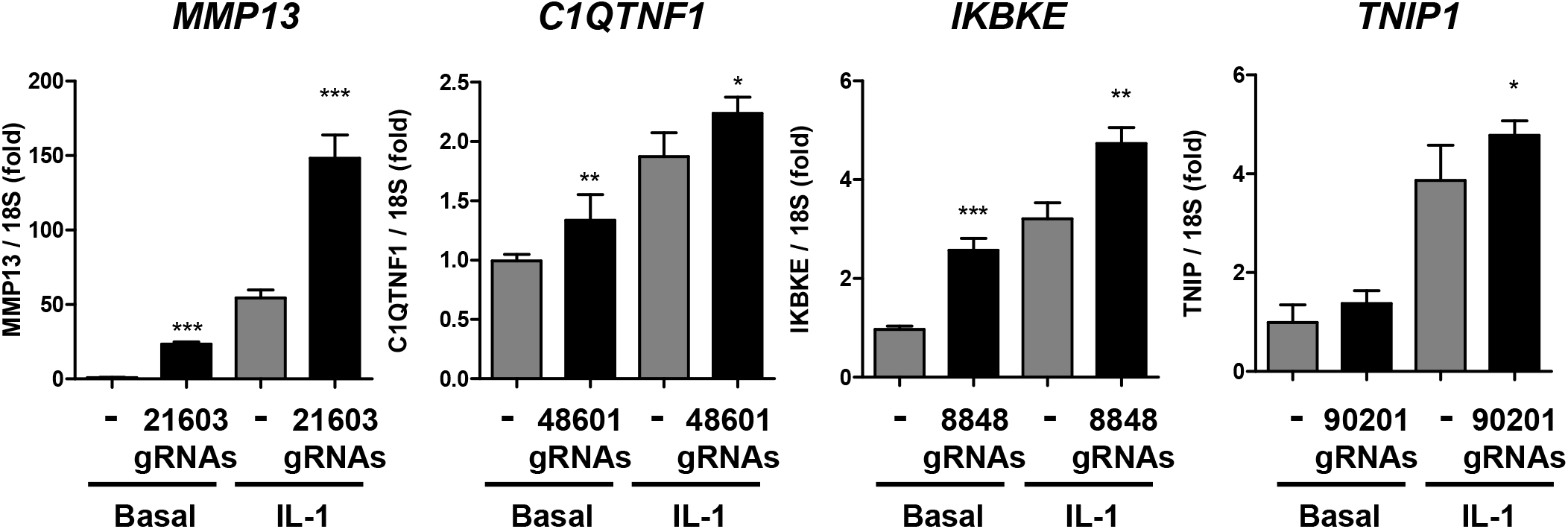
Effect of enhancer-targetted Cas9-VPR on gene expression. Expression of the indicated genes measured by real-time qRT-PCR in SW1353-Cas9-VPR cells following transfection of ATAC-seq region gRNAs and IL-1 stimulation for 6 hours. Numbers 21603, 48601, 8848, 90201 refer to the ATAC-seq peak identifier defined by MACs. Values are mean ± SEM of data pooled from two independent experiments (performed in sextuplate). *P<0.05; **P<0.01; ***P<0.001 for deletion versus wild-type for each control or IL-1 treatment group. Significant differences between sample groups were assessed by a two-tailed Student’s t-test.

Finally, to further assess the ability of these regions to act as IL-1-responsive enhancers the regions were cloned into an enhancer luciferase reporter and the impact of stimulation determined. The regions remaining following the Cas9 deletion were cloned as controls. The accessible regions at the *C1QTNF1* and *IKBKE* loci promoted luciferase expression following IL-1 stimulation for 6 or 24 hours (Figure 5), while for *MMP13* region only the later time point saw an increase in luciferase activity. The region within *TNIP1* did not significantly drive luciferase expression at either time. Neither the *MMP13*nor *C1QTNF1* Cas9-deleted regions were able to drive IL-1 induction of luciferase activity but the *IKBKE* deletion region did show some residual low level ability to enhance luciferase expression at 24 hours post IL-1 treatment.

**Figure 5.**
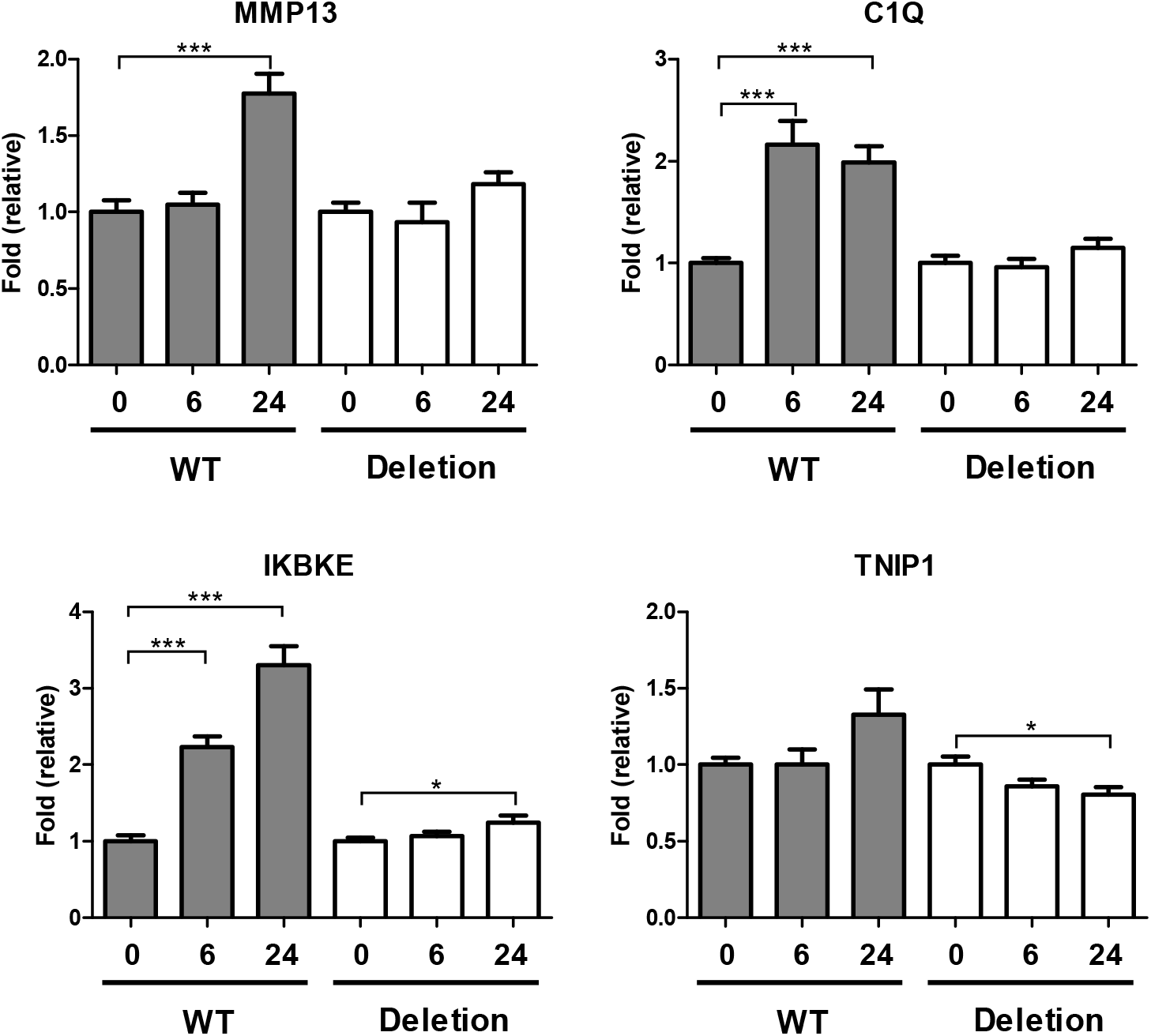
Enhancer activity of differentially accessible chromatin regions. Luciferase activity in SW1353 cells transfected with pGL3-promoter containing wild-type and deleted ATAC-seq peak containing regions following IL-1 stimulation for 6 or 24 hours. Values are mean ± SEM of data pooled from two independent experiments (each performed in sextuplate). *P<0.05; ***P<0.001 for IL-1 treatment versus unstimulated cells. All firefly luciferase data were normalised to Renilla levels. Significant differences between sample groups were assessed by one-way analysis of variance followed by the Bonferroni post-hoc test.

## Discussion

Herein we established the chromatin accessibility landscape in the well-characterised IL-1 responsive SW1353 chondrocyte cell line following IL-1 stimulation [25]. We identified 241 regions exhibiting changes in chromatin accessibility, a large proportion of which correspond with enhancer regions of chondrocytes. Importantly, we have experimentally examined the function of a number of these differentially accessible regions and confirmed their enhancer activity and contribution to IL-1-induced gene expression.

Changes in chromatin conformation and accessibility are widespread during development where cells undergoing differentiation exhibit major alterations in gene expression [1]. Consistent with this, comparison of chromatin differences between cell types by ATAC-seq has identified large numbers of regions of differing accessibility [28], supporting previous findings, using the similar DNase I sensitivity assay, that chromatin accessibility at expression quantitative trait loci (eQTLs) is the major determinant of human expression variation [29]. Many such accessible regions overlap with enhancer modifications confirming the cell type specific regulation of gene expression by enhancers. The impact of signalling pathway activation on chromatin accessibility is less well established. Calderon *et al*. used distinct stimuli to induce biologically relevant responses in a range of T and B cell subsets and monocytes and NK cells and assessed chromatin accessibility via ATAC-seq [30]. Overall they found dramatic changes in the chromatin landscapes of the B and T cells, but limited effects in innate lineage cells. Macrophages, differentiated from induced pluripotent stem cell (iPSC) lines, have also been assessed for stimuli-dependent chromatin accessibility changes, including using IFNγ [31]. Similarly, murine macrophage enhancers have been classified into several categories based on their response to a number of distinct stimuli. This identified a relatively small number (2140) of genomic regions that were unmarked (lacking H3K4me1 and H3K27Ac histone modifications) in unstimulated cells but which acquired enhancer features upon at least one stimulation, including IL-1β, which were termed latent enhancers [32]. We observed our differentially accessible chromatin regions overwhelmingly corresponded to regions marked by H3K4me1 and H3K27Ac in MSC-derived chondrocytes, thereby categorised as enhancers. Without chromatin state data from control or IL-1-stimulated SW1353 cells we can only speculate whether our differentially assessable regions have histone marks characteristic of enhancers in the absence or presence of IL-1 stimulation, in contrast to latent enhancers. Recently, Weiterer *et al*. determined IL-1α responsive enhancers in HeLa-derived KB cells using ATAC-seq [23]. In contrast to our data, they identified >75,000 IL-1-induced open regions although their experiment was only in singlicate and therefore they could not determine significance. However, when commonly co-analysing the two datasets it was apparent regions becoming more accessible following IL-1 treatment significantly overlapped when compared to those becoming less accessible, perhaps unsurprisingly suggesting important inflammatory regulated regions of the genome are cell-type independent.

De novo motif discovery within the latent enhancers defined following TNFα or IL-1β stimulation of macrophages identified significant RelA/NF-κB over-representation [32]. Similarly, NF-κB motifs are enriched in accessible regions also marked by H3K27Ac following IL-1αstimulation [23]. Within the IL-1 increased accessible regions in our dataset a similar, highly significant, enrichment for RelA/NF-κB binding motifs was observed. Analysis of the GTRD ChIP-seq database and IL-1-induced RelA ChIP-seq data confirmed that the four regions focussed upon were also bound by RelA in cells. While NF-κB is not thought to be a ‘pioneering factor’, i.e. a transcription factor able to initiate chromatin remodelling, evidence does support a role for the factor in changing the nucleosome landscape at activated loci [33, 34]. The second most enriched motif in IL-1 increased accessible regions was IRF1, interferon regulatory factor 1, a factor that increases with IL-1 stimulation [35]. We also identified enrichment for the ELF3 binding motif, which again is induced by IL-1 and is a crucial mediator of the phenotypic and transcriptional changes induced by this cytokine, including in chondrocytes [36, 37], where it can also regulate *MMP13* expression [38]. Chondrocyte-specific *Elf3* deletion also reduces the severity of disease in a murine osteoarthritic model [39]. Thus, our analysis identifies physiologically relevant additional IL-1-responsive factors besides NF-κB, though we cannot rule out that the activation of these transcription factors are secondary events due to our chosen 6 hour time point. Motifs enriched in the less accessible peak regions were more diverse but did not contain RelA/NF-κB. However, enriched motifs included for factors known to also be regulated by IL-1, or to co-ordinate with NF-κB, including IRF1 again, RUNX and STAT3/6 [40, 41].

IL-1 treatment of cells predominantly activates NF-κB and MAPK signalling leading to the transcriptional regulation of a wide variety of genes. However, the role of chromatin alterations and enhancers in this activation is largely unknown. Stimulation of SW1353 with IL-1 for 6 hours results in expression changes of greater than 1.5-fold in almost 500 genes. It is unclear why only a small proportion of IL-1-induced genes exhibited changes in chromatin accessibility but where an accessible region was linked to an IL-1-regulated gene there was a strong positive correlation, suggesting that accessibility influences gene expression.

We confirmed the function of a number of accessible regions in regulating IL-1-induced gene expression, focussing on regions that became more accessible with stimulation. We chose to focus on regions within *MMP13*, *C1QTNF1*, *IKBKE* and *TNIP1* because all had strong evidence for the binding of RelA following IL-1 stimulation. These accessible regions were all intronic to the purported regulated gene, and the deletion of these regions within *MMP13*, *C1QTNF1* and *IKBKE* lead to a reduction in the respective gene expression following IL-1 stimulation, highlighting intronic enhancers and their role in gene regulation. However, none of the regions deleted completely ablated the induction of the corresponding gene, and in fact deletion of region 90201 near *TNIP1* had no effect on the genes expression. These data are consistent with a model of multiple enhancers looping to control expression and suggests an important role for enhancer redundancy in the regulation of inflammatory signalling, as has been seen during development where the purported mechanistic function is to prevent deleterious phenotypic consequences [42]. No impact of intron deletion was observed on the basal level of gene expression suggesting splicing was not impacted. Although all four regions were activated using a dCas9-VPR transactivation system, when using an enhancer-reporter construct, again only the *TNIP1* region failed to show enhancer-like activity.

MMP13, matrix metalloproteinase-13, is a collagenase responsible for cartilage collagen turnover and pathological destruction in osteoarthritis progression. MMP13 is highly regulated by inflammatory signalling, the inhibition of which is a proposed therapeutic strategy in OA, however the mechanism by which cytokines directly impact on its expression has proven to be somewhat elusive [43]. Proximal promoter AP-1 factor binding is critical for the accurate inflammatory induced expression of the gene and in part the binding of ATF3, however, NF-κB is also critical for activating cytokine-induced *MMP13* expression [44–47]. The data here suggests this could in part be via a novel RelA-bound intron 5 enhancer. Two distal enhancers at −10kb and −30kb in osteoblasts may also contribute to basal and Vitamin D receptor, or RUNX2-mediated, *MMP13* expression [48]. IKBKE, TNIP1 and C1QTNF1 are themselves important molecules in inflammatory signalling [49–53] and known to be regulated in an NF-κB-dependent manner [52, 54].

In conclusion, we have identified a number of dynamic chromatin accessibility changes modulated by short stimulation with IL-1. Regions that become accessible are enriched for RelA binding motifs and chromatin enhancer marks. Finally, we have functionally validated the regulatory function of several regions using a range of tools, including CRISPR/Cas9-mediated genome editing. Of particular interest our data suggests the presence of an intronic enhancer important for the regulation of MMP13 following cytokine stimulation. Our data confirms that ATAC-seq represents a simple method to identify novel and highly dynamic chromatin changes and potentially important enhancer regions following cell stimulation.

## Acknowledgements

This work was supported by the Medical Research Council and Arthritis Research UK as part of the MRC-Arthritis Research UK Centre for Integrated Research into Musculoskeletal Ageing (CIMA, grant references JXR 10641 and MR/P020941/1); the Newcastle upon Tyne Hospitals NHS Charity; the JGW Patterson Foundation; and The Dunhill Medical Trust (R476/0516).

## Declaration of contributions

DAY and MJB were involved in concept and study design. JF, ACP, and MJB were involved in data collection. KC, MJB, LNR and DAY were involved in data analysis and interpretation. DAY, MJB and DJD obtained the funding. All authors were involved in drafting the manuscript and gave final approval of the version submitted.

---

**Supplementary Figure 1**

**A.** ATAC-seq sequencing fragment size distribution for pooled data from Control (n=3) and IL1 (n=3) sample libraries. The banding pattern is indicative of nucleosome free regions, mono-, and dinucleosomes. **B.** nucleoATAC signal aggregated over all genes indicating nucleosome positioning around transcription start sites (TSS). Positive and negative numbers indicate nucleosomes upstream and downstream of the TSS.

**Supplementary Figure 2**

Distribution of all the ATAC-seq peaks with regards to Genbank genic annotation as defined using HOMER [14].

**Supplementary Figure 3**

Comparison of IL-1 (**A**) and basal/control (**B**) ATAC-seq peaks from this study with Weiterer *et al*. [23]. Using identical analysis tools there is little overlap in overall identified peaks (control or IL-1 enriched. Both datasets were handled as though performed in singulate and, following MACS2 analysis, peaks 2 enriched over the comparator counted (i.e. increased 2-fold in IL-1 vs. control). Although there was little overlap in the overall peaks in the two datasets >50% of our significant IL-1-induced open regions were IL-1 peaks in the Weiterer *et al*. data (**A**) while there was no enrichment for less accessible peaks following IL-1 stimulation in the control group (**B**).

**Supplementary Figure 4**

Extension of Figure 2B a comparison of significant ATAC-seq peak accessibility fold change with expression change in proximal labelled gene. Green and red represent genes significantly up or down regulated respectively following IL-1 stimulation.

**Supplementary Figure 5**

Genomic vignettes of the four putative enhancer regions examined with included MACS2 RelA ChIP-seq data from GTRD.

**Supplementary Figure 6**

IGV and UCSC genome browser generated vignettes of JEME-derived interaction predictions which indicate that the IL-1 induced enhancers (*) for MMP13, C1QTNF1 and TNIP1 would interact with the promoters/TSS of the corresponding gene.

**Supplementary Figure 7**

Real-time qRT-PCR of IL-8 expression (relative to 18S) in control (shaded) and cell-lines deletion for the indicated putative enhancers following 6 hrs of IL-1 stimulation. Data confirms that deletion of the enhancers examined did not affect NF-κB signalling *per se*.

