## Supplemental Figures for "Dynamic chromatin accessibility landscape changes following interleukin-1 stimulation"

### Supplemental Figure 1

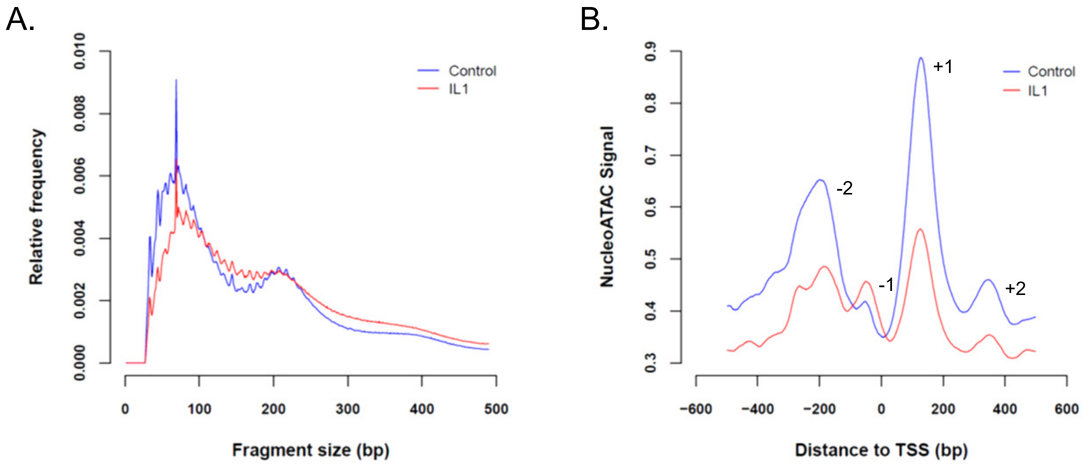


### Supplemental Figure 2 – Genbank annotation of peak distribution

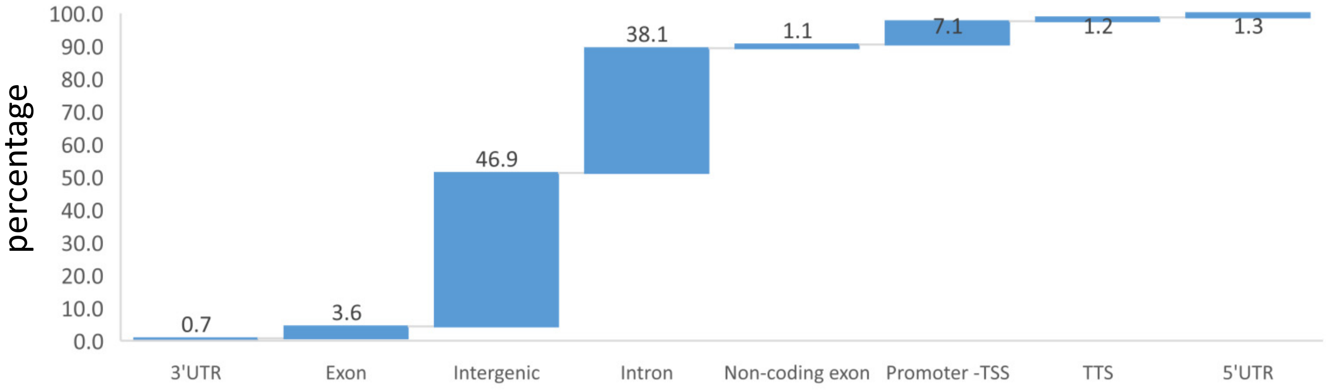


### Supplementary Figure 3

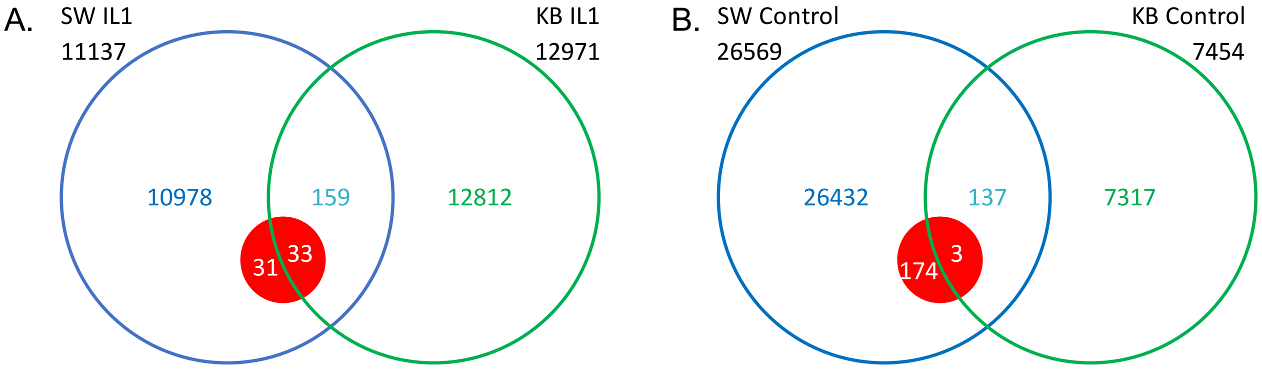

### Supplemental Figure 4

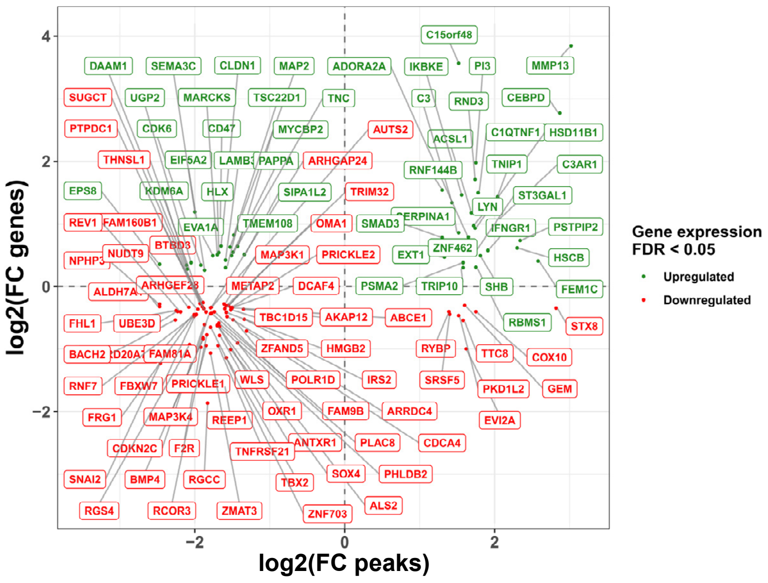


### Supplementary Figure 5

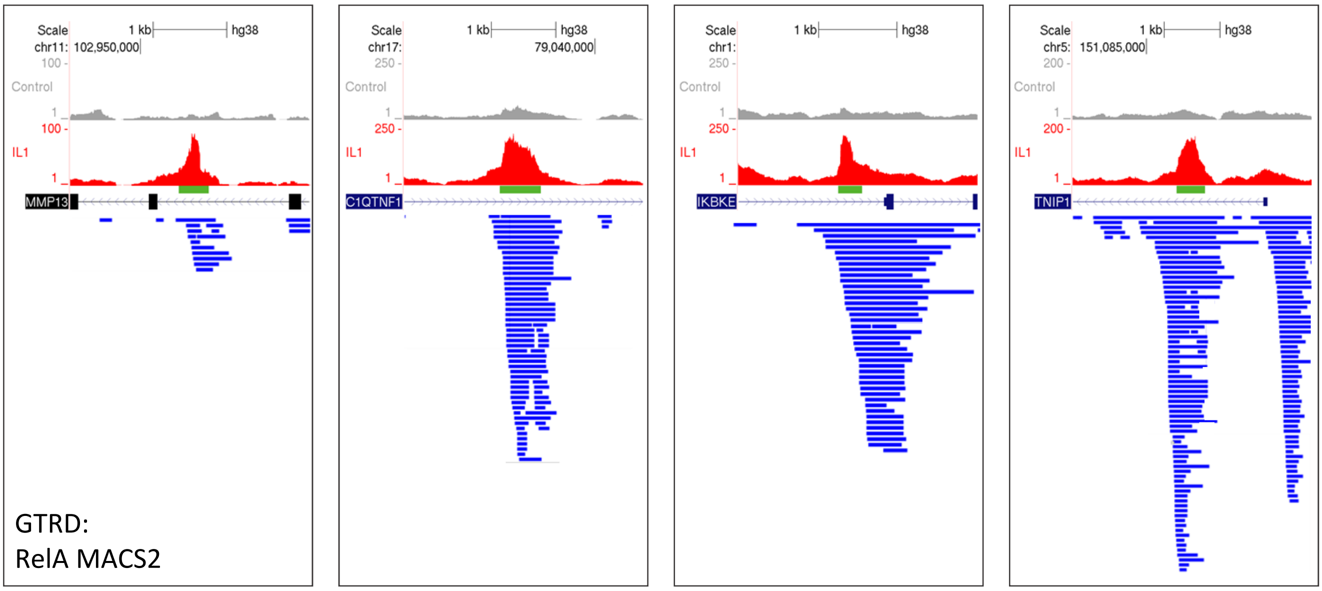

#### Supplementary Figure 6

##### MMP13

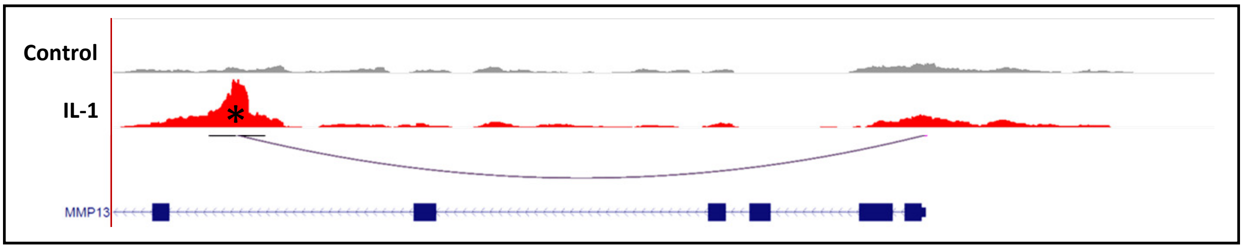


##### C1QTNF1

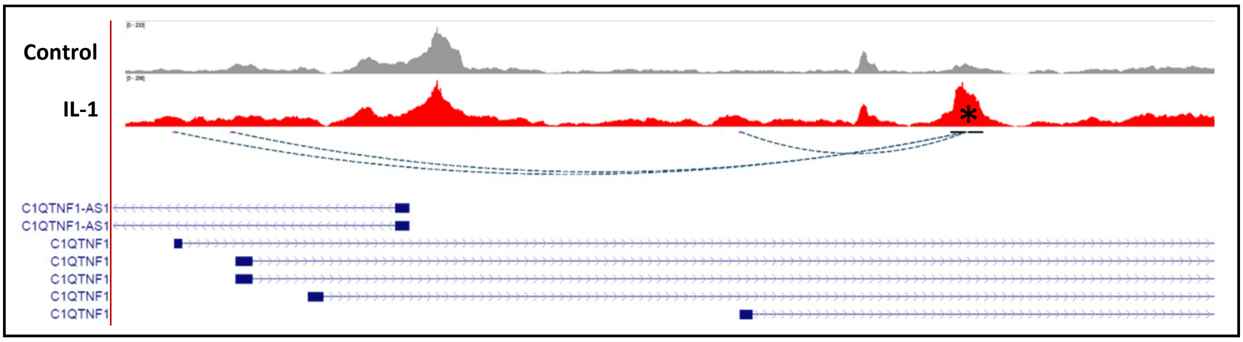


##### TNIP1

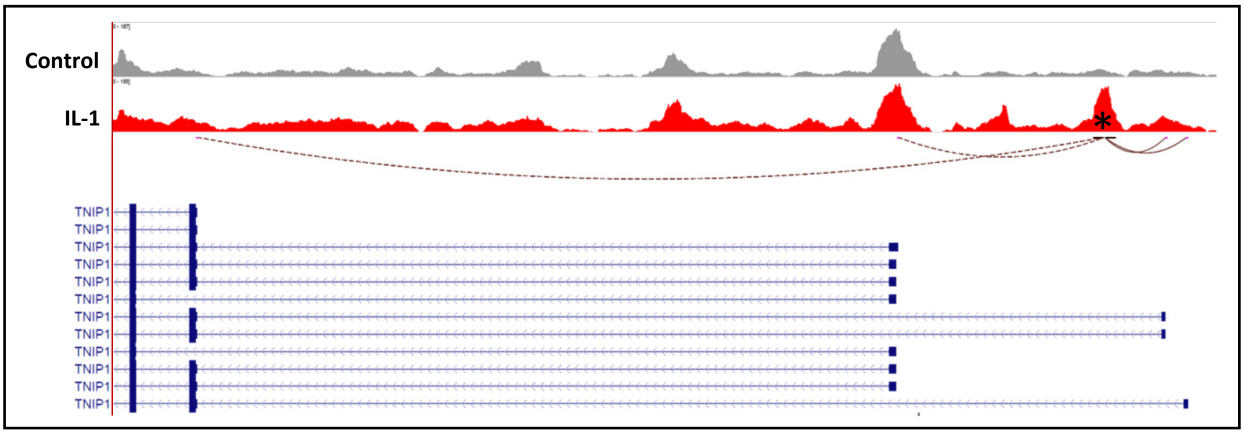


#### Supplemental Figure 7

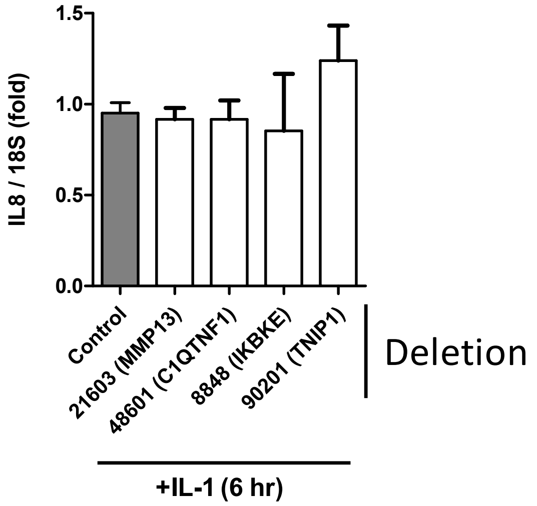
